# A Multi-Omics Framework Reveals Phosphorylation-Dependent Control of Protein Interactions

**DOI:** 10.64898/2026.07.14.738447

**Authors:** Kosuke Ogata, Emmanuel Matabaro, Ellen Aarts, Jürgen Jänes, Yasushi Ishihama, Pedro Beltrao

## Abstract

Protein phosphorylation regulates nearly every cellular process, yet most of the hundreds of thousands of human phosphosites remain functionally uncharacterized. Rather than prioritising phosphosites by conservation or structural features, here we use the abundance of an interaction partner as a readout of whether a phosphosite affects that interaction. This idea exploits the fact that subunits of stable complexes are often degraded when unbound. Here, we apply a nested linear regression model to pan-cancer data from 1,006 tumours, while controlling for transcriptional and other covariates. We identified 6,160 associations between 3,038 phosphosites and the abundance of interacting proteins, including several known interaction-regulating sites. Mapping these onto AlphaFold-predicted complexes placed 239 sites at interaction interfaces, while another 402 were linked to compartment-specific localisation, indicating that phosphorylation can also tune interactions by relocating proteins between compartments. Affinity-purification mass spectrometry of NKAP and NUF2 phosphosite mutants experimentally supported some of these predictions. Together, this framework reveals a widespread coupling between phosphorylation and interaction-dependent protein abundance and provides a prioritized, structure-informed resource for characterizing the human phosphoproteome

## Introduction

Protein phosphorylation serves as a molecular switch that regulates virtually every cellular process, from signal transduction and metabolism to cell division and cell death (“Signaling—2000 and Beyond” 2000). This reversible modification represents one of the most prevalent post-translational modifications in eukaryotic cells, with most proteins undergoing phosphorylation during their lifecycle (Hornbeck et al. 2015). By altering protein stability, conformation, activity, interaction affinities to other molecules, and subcellular localizations, phosphorylation orchestrates the dynamic protein interaction networks that govern cellular responses to environmental stimuli (Humphrey et al. 2015).

Despite phosphorylation’s central role in cellular regulation, determining which phosphorylation events are crucial for functional consequences remains a fundamental challenge (Needham et al. 2019; Ochoa et al. 2020a). The rise of phosphoproteomics technologies led to the identification of hundreds of thousands of phosphosites in the human proteome, yet many lack functional characterization (Sharma et al. 2014; Ramasamy et al. 2020; Viéitez et al. 2022). Mutational studies have been used to rank the importance of phosphorylation for cellular fitness, but cannot yet be applied to the human phosphoproteome at scale (Viéitez et al. 2022; Schastnaya et al. 2021; Kennedy et al. 2024). Beyond studying the importance of phosphorylation on cellular fitness, experimental approaches have been developed to study the role of phosphorylation in regulating for example metabolism (Oliveira et al. 2015; Schastnaya et al. 2021), protein stability (Potel et al. 2021; Smith et al. 2021) or protein solubility (Sridharan et al. 2022). Computational studies have sought to differentiate between functional and nonfunctional phosphorylation sites by examining their evolutionary conservation (Boekhorst et al. 2008; Beltrao et al. 2012), kinase specificity (Landry et al. 2009), and spatial arrangement within protein-protein interaction interfaces (Nishi et al. 2011; Betts et al. 2017; Šoštarić et al. 2018), or combinations of such features (Torres et al. 2016; Beltrao et al. 2012; Ochoa et al. 2020a).

We propose here an alternative approach to predict phosphosites that modulate interactions by leveraging protein co-abundance patterns. Protein co-abundance has been often used to establish ’protein-protein associations’, a term that we use here to include direct physical interactions but also other forms of co-abundance driven protein associations (Gonçalves et al. 2017; Sousa et al. 2019; Laman Trip et al. 2025). The principle of protein co-abundance offers a useful framework for identifying functional regulatory relationships, including those mediated by phosphorylation (Sousa et al. 2019). This approach builds on the observation that, for stable protein complexes, their subunits are often co-expressed with excess unpartnered proteins typically degraded to preserve cellular homeostasis (Gonçalves et al. 2017; Sousa et al. 2019). Co-abundance has been shown to be a simple but accurate predictor of protein-protein associations because of this stoichiometric coupling (Laman Trip et al. 2025). We have previously shown that, by incorporating transcriptomic data as a confounding variable, co-abundance analysis can isolate post-transcriptional regulatory effects, showing how the presence or modification of one protein can directly affect its partners’ abundance (Gonçalves et al. 2017). We have previously shown, with limited data, that when applied to phosphorylation data, this can identify regulatory phosphosites whose modification state correlates with partner protein levels, effectively mapping the phosphorylation events that control complex formation (Sousa et al. 2019).

Recent technological advances have transformed our ability to apply co-abundance analysis at scale while addressing the historical lack of structural context. Large-scale proteomic and phosphoproteomic datasets, such as those from the Clinical Proteomic Tumor Analysis Consortium (CPTAC) (Li et al. 2023), now provide the higher statistical power needed to detect such regulatory relationships at larger scale. At the same time, AlphaFold modelling now enables mapping of phosphosites onto predicted structures. This combination of increased sample size for proteomic data and structural predictions creates an opportunity to contextualize phosphorylation-mediated regulation within its structural environment.

Here we apply this co-abundance-based framework at proteome scale, identifying thousands of phosphosites whose modification state tracks the abundance of an interaction partner. Mapping these phosphosites onto AlphaFold-predicted complexes resolves those that sit at interaction interfaces. Integrating phosphosite-resolved subcellular localisation data identifies a second class that acts by relocating proteins between compartments. Finally, affinity-purification mass spectrometry of selected baits and phospho-mutants confirms that individual phosphosites reshape specific interactions. The result is both a mechanistic account of how phosphorylation tunes interaction-dependent protein stability and a structure-informed, prioritised resource for experimental follow-up across the phosphoproteome.

## Results

### A co-abundance model identifies phosphosites that control the abundance of interaction partners

We set out to identify phosphosites whose modification alters a protein-protein interaction, using the abundance of the partner protein as the functional readout (Figure 1A). We constructed a comprehensive dataset from CPTAC pan-cancer data encompassing 1,006 samples (Figure 1A-C). The dataset includes gene copy number variation (CNV) measurements, transcript abundances, proteome quantification, phosphoproteome profiles, and associated metadata (age, sex, and cancer type). To ensure robust association analyses, we retained proteins and phosphosites quantified in more than 200 samples. All log2 intensities across transcriptome, proteome, and phosphoproteome datasets were normalized to minimize batch effects. The proteome data consisted of 11,231 proteins, while phosphoproteome data consisted of 6,635 proteins, leaving an overlap of 6,073 proteins (Figure 1D).

**Figure 1.**
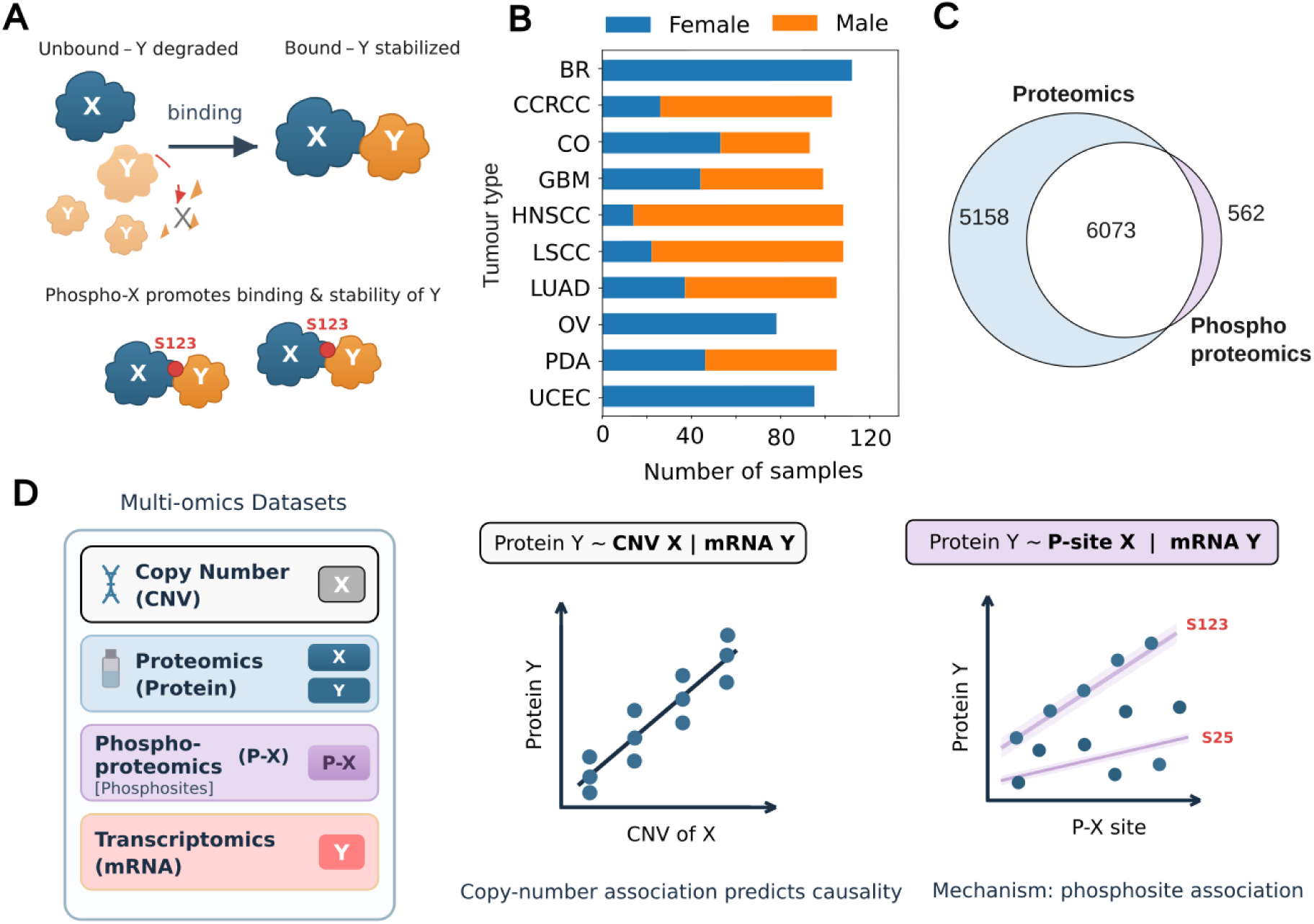
Compendium of multiomics data to study phosphorylation control of protein interactions. (A) Schematic of the modeling approach used to test associations between protein X and protein Y. (B) Workflow overview of the CPTAC pan-cancer dataset curation, integrating CNV, transcriptomics, proteomics, phosphoproteomics, and clinical metadata. (C) Distribution of samples across the cohorts. (D) Venn diagram illustrating the overlap between the proteome (11,231 proteins) and phosphoproteome (6,635 proteins) datasets, resulting in 6,073 common proteins.

Using the curated dataset, we performed systematic linear regression analyses on protein pairs restricted to documented protein interaction partners, as cataloged in established protein interaction databases, such as BioGRID and CORUM (see Methods). We first tested if the copy-number state (CNV) of protein X, is associated with a change in protein levels of its interactor Y when controlling for any change in RNA levels of Y. This is first done at copy-number level because this provides a sense of causal directionality. Because the copy number of gene X is set upstream of protein abundance and is not itself influenced by the level of protein Y or by downstream cellular state. An association between the CNV of X and the protein level of Y is more readily interpreted as X regulating Y than the converse. We identified 28,444 CNV-protein with significant associations (FDR < 0.01).

We ran a second association model between the transcript level of X and the protein level of the interacting protein Y, when controlling for its transcriptional changes of Y, yielded substantially more significant pairs (n = 396,156). This very large number of associations is likely to contain many spurious indirect associations. We retained only associations where CNV and transcript effects on target interacting protein abundance were significant and directionally consistent, resulting in 12,224 protein-protein pairs whereby the levels of protein X is likely to regulate the abundance of its interacting protein Y (Figure 2A).

**Figure 2.**
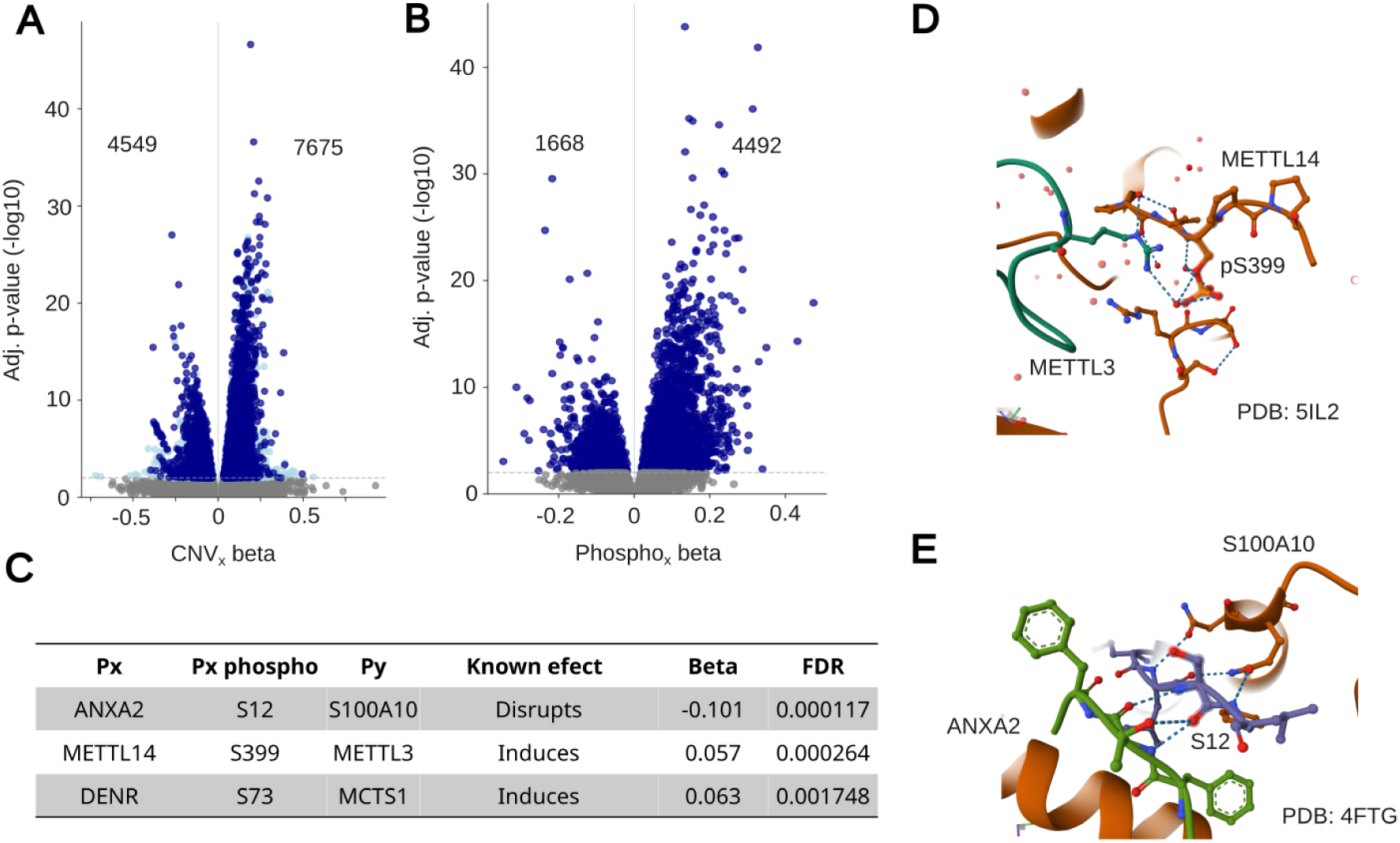
A co-abundance model identifies phosphosites that control the abundance of interaction partners. (A) Volcano plot of the protein-protein pairs with significant CNV associations, highlighted in dark blue are the 12,224 with directionally consistent CNV and transcript effects. (B) Volcano plot of the phosphosite-protein pairs with significant associations with changes in abundance of interacting proteins. (C) Examples of known interaction-regulating sites, including examples with known structural rational, such as (D) METTL14 S399 (positive association with METTL3 abundance) and (E) ANXA2 S12 (negative association with S100A10).

For these 12,224 protein interactions we next examined if the phosphorylation levels of protein X would provide a better explanation for the protein levels of protein Y, indicative of a phosphorylation-mediated control of protein interactions (**see Methods**). From 46,478 initial phosphosite-protein pairs, we identified 3,038 phosphosites predicted to regulate 6,160 interactions (Figure 2B). These include phosphosites with established roles in regulating protein interactions (Figure 2C-E). For example, METTL14 S399 phosphorylation, which promotes METTL3-METTL14 interaction at the protein interface, exhibited a positive association with METTL3 abundance, consistent with phosphorylation-enhanced complex stability (Figure 2C and 2D). Similarly, ANXA2 S12, known to disrupt the ANXA2-S100A10 interaction, showed a significant negative association with S100A10 protein abundance in our analysis, consistent with what is reported in the literature (Figure 2C and 2E). These examples demonstrate that our co-abundance approach successfully captures known phosphorylation-mediated regulatory mechanisms.

### Localization-regulating phosphosites couple subcellular compartment to partner abundance

We next investigated potential mechanisms by which the phosphorylation levels could impact on protein interactions. We first looked at phosphosites that potentially regulate subcellular protein localization or exhibit distinct subcellular distribution patterns compared to their parent proteins. To identify localization-regulatory phosphosites, we constructed a catalog of localization related phosphosites using two complementary approaches. First, we extracted phosphosites annotated with “intracellular localization” functions from the PhosphoSitePlus database, which curates experimentally validated phosphorylation sites with known regulatory roles. Second, we compiled phosphosites displaying distinct subcellular distribution patterns from published subcellular phosphoproteomics studies, identifying sites where phosphorylation status correlates with specific cellular compartments. This analysis revealed that 402 of our 3,038 significantly associated phosphosites (13.2%) possessed documented or predicted roles in subcellular localization regulation (Figure 3A). Notably, these phosphosites tend to have higher predicted functional scores (Figure 3B), indicating that these 402 phosphosites are more likely to be functional based on a previous machine learning model that integrated diverse features such as conservation, degree of regulation, among others (Ochoa et al. 2020b). To further evaluate these phosphosites, we examined the original subcellular fractionation data from Martinez-Val et al. (Martinez-Val et al. 2021), comparing the compartmental distribution profiles of phosphorylated proteins with those of their associated target proteins. We observed several phosphosites where the modified protein exhibited subcellular distribution patterns more like the interacting protein Y than to the unmodified protein X (Figure 3C-F). An example is LRWD1, also known as ORCA, a protein known to recruit and stabilize the origin recognition complex (ORC) onto chromatin during G1 (Shen et al. 2010). Our analysis predicts that phosphorylation of LRWD1 at S212 promotes binding to ORC subunits (ORC2 and ORC3, Figure 3D). The compartmental distribution data fits this model with the S212 phosphorylated form of LRWD1 having a distribution that is more correlated with ORC2/3 than the non-phosphorylated form of the protein (Figure 3D). These examples illustrate how spatial regulation can represent a significant mechanism by which phosphorylation sites influence protein-protein interaction networks.

**Figure 3.**
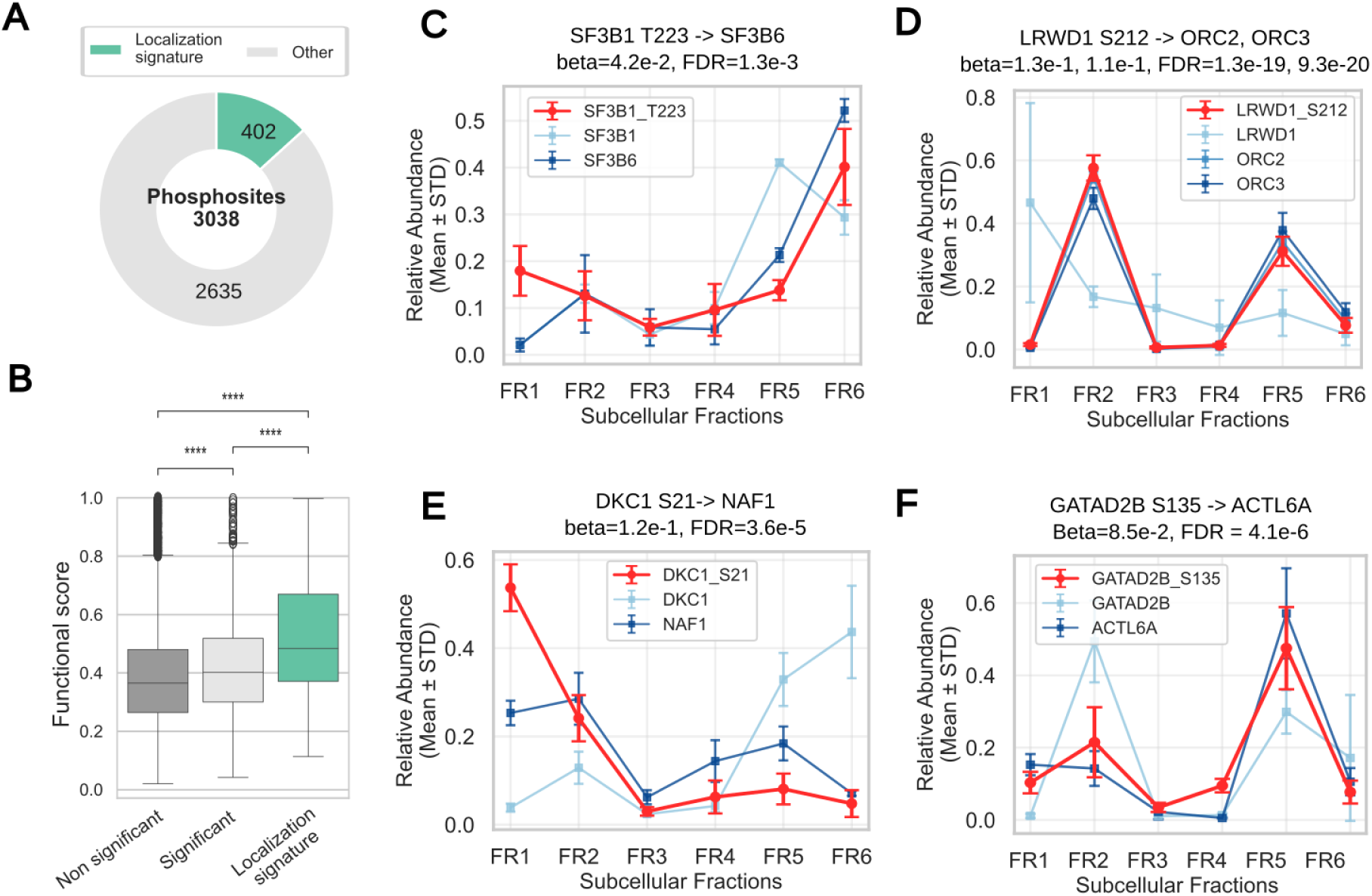
Localization-regulating phosphosites couple subcellular compartments to partner abundance. (A) Proportion of the 3,038 significantly associated phosphosites with documented or predicted roles in subcellular localization regulation (n = 402). (B) Comparison of predicted functional scores between localization-regulating phosphosites and other significant sites. (C-F) Subcellular fractionation profiles comparing the compartmental distribution of phosphorylated proteins to their interaction partners.

### Phosphorylation Sites Located at Protein Complex Interfaces

If a phosphosite controls a partner’s abundance by modulation of interactions, it should frequently sit at physical interfaces between proteins, either at the direct interface of the modulated interaction or at the interface with a third-protein (Figure 4A). Given the limited availability of experimental protein complex structures we turned to AlphaFold predicted structures of protein complexes. We first used AlphaFold-Multimer predictions from a previous study (Jänes et al. 2024) (486,099 structures), covering 1,385 of the 3,211 protein pairs of our interest. For the remaining pairs, we tried to generate additional structures using AlphaFold (ColabFold, see Methods), successfully obtaining models for another 1,410 pairs, achieving coverage of 2,795 of 3,211 total pairs (87.0%). To ensure structural reliability, we applied quality filters based on predicted DockQ (pDockQ) scores, retaining only models with pDockQ > 0.23 for interface analysis, leaving 847 medium to high-quality structures (Figure 4B).

**Figure 4.**
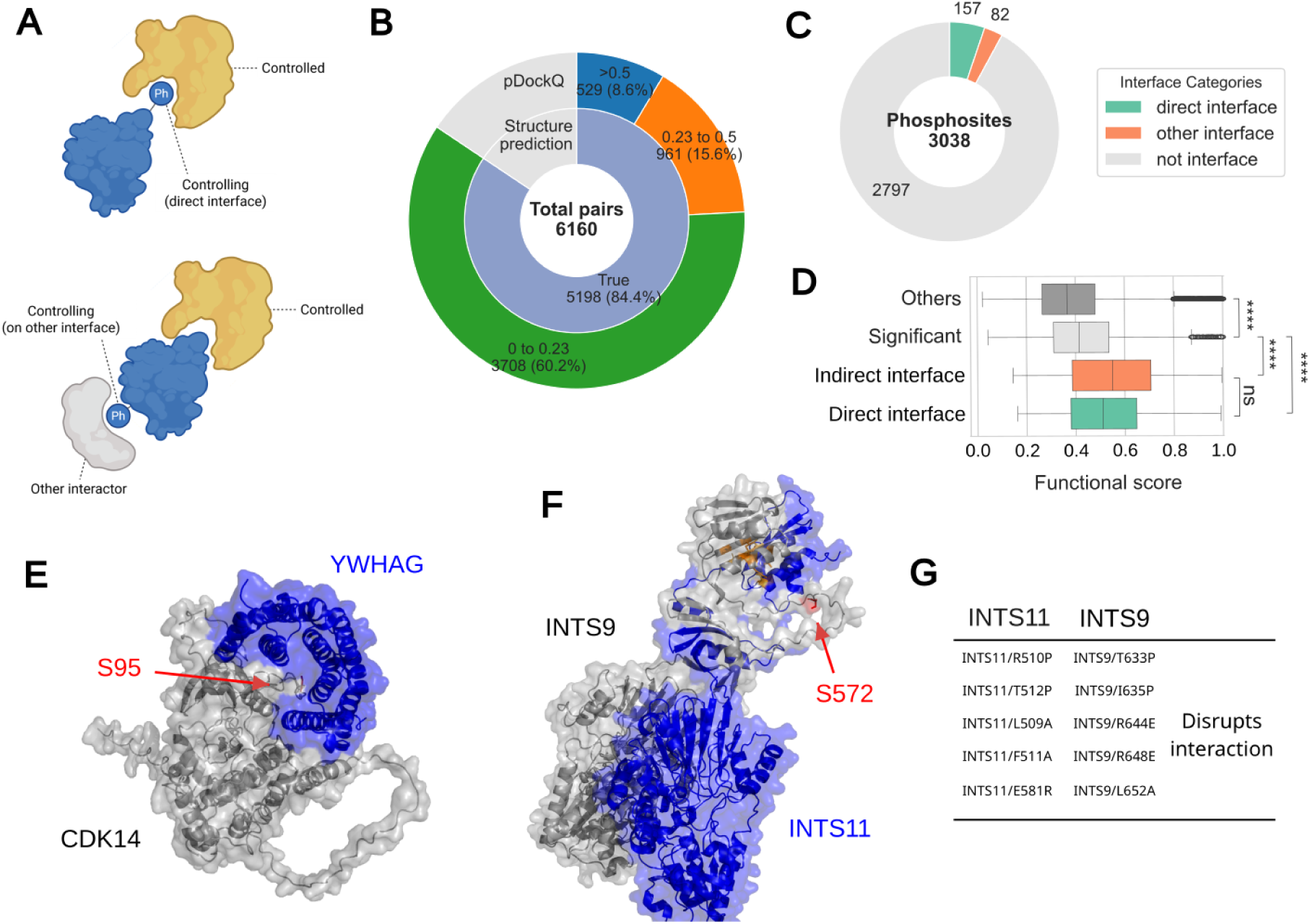
Phosphorylation sites predicted to regulate interactions located at protein interfaces. (A) Schematic representation of phosphosites positioned at physical protein-protein interaction interfaces either directed at the interface of modulated partners or at the interface of a third protein. (B) We attempted to model 6,160 phosphosites-interaction pairs, including multiple phosphosites per structural model of a complex. Of these, 5,198 could be modelled by AlphaFold with, 961 found in medium-quality and 529 in high-quality structures based on pDockQ. (C) Visualization of the 239 phosphosites identified at protein complex interfaces. (D) Comparison of functional scores between interface-located and non-interface phosphosites. (E) Example showing CDK14 phosphorylation enhancing YWHAG binding. (F) Example showing INTS9 S349 phosphorylation potentially strengthening the INTS9-INTS11 interaction and (G) known mutations known to disrupt this interaction at residues near the phosphosite position.

Interface residues were defined as amino acids in close proximity to the interacting partner, with distances calculated between any atoms of a given residue and the nearest atoms in the partner protein (Figure 4A). We evaluated different distance cutoff criteria (5, 8 and 10 Å) and a filter based on the predicted local distance difference test (pLDDT). Phosphorylation predominantly occurs within intrinsically disordered regions, which typically exhibit lower confidence scores in structural predictions. Applying a stringent pLDDT cutoff of >70 would eliminate more than half of interface phosphosites with each distance criterion applied, potentially excluding functionally relevant sites (Figure S1A). The pLDDT cutoff even excluded the METTL14 S399 and ANXA2 S12, which have been shown to be located at the interface in experimental complex structures (Figure 2D,E). Therefore, we retained interface phosphorylation sites regardless of pLDDT values to maximize coverage of potentially regulatory events. Also, the 8 Å distance criterion successfully recovered both METTL14 S399 and ANXA2 S12 as interface phosphosites, while the 5 Å criterion excluded METTL14 S399. Thus, for defining interface and near-interface phosphosites, we accepted sites with less than 8 Å from interactors as interface sites (Figure S1A). We note that 8 Å will contain some fraction of sites that are not in direct contact but will be very close and therefore could still influence the interaction.

We also investigated phosphosites located at interfaces of other protein complexes involving the same proteins (Figure 4A). Such phosphosites may contribute to stable complex formation with other proteins and could indirectly or allosterically influence the target interaction through conformational changes, competitive binding or cellular re-localization. To annotate these interface residues, we used all the binary complex structure prediction data. Here, the amino acid residue on a specific protein is considered as interface residue if it falls in the interface category in any of binary complexes. To prevent over-annotation of interface residues, we applied more stringent criteria for this extended analysis: pDockQ > 0.5, interface distance < 5 Å, and pLDDT > 70. These criteria yielded a reasonable fraction of residues classified as interface-associated while maintaining structural confidence (Figure S1B). Our structural analysis identified 239 phosphosites located at protein complex interfaces (Figure 4C).

Like for phosphosites controlling localisation, these interface-located phosphosites exhibited higher predicted functional scores compared to non-interface sites, suggesting greater functional importance (Figure 4D). Several examples illustrate the biological relevance of our findings. CDK14 phosphorylation at a site classified as interface-located showed a positive association with YWHAG abundance. YWHAG (14-3-3 gamma) is a well-characterized phospho-binding protein recognizing specific phosphorylated motifs. The CDK14 phosphosite matches the canonical 14-3-3 binding motif, indicating that phosphorylation likely enhances the CDK14-YWHAG interaction (Figure 4E). Another example involves the INTS9-INTS11 interaction, where phosphorylation showed significant association with target protein abundance. This interaction is sensitive to interface mutations according to the IntAct mutation database, in part validating the predicted structure, and our analysis suggests that INTS9 S349 phosphorylation may strengthen the interaction between these integrator complex subunits (Figure 4F). These examples provide mechanistic support for our model that interface-located phosphorylation sites can regulate protein-protein interactions and thereby modulate target protein stability.

### Affinity-purification mass spectrometry confirms site-specific interaction changes

To test directly whether the predicted regulatory phosphosites alter interactions, rather than merely co-vary with partner abundance, we performed AP-MS on three baits, NKAP, METTL14, and NUF2, each as wild-type and as candidate phosphosite mutants, that we predict could impact on interactions with their cognate interaction partners DDX41, METTL3 and NDC80 respectively. We selected 3 phosphosites in NKAP (S149, S157 and S161), and one for METTL3 (S399) and NUF2 (S247) and we mutated each to alanine (phospho-null) or glutamate as a phospho-mimetic mutation. We included METTL3 S399 as a previously described example for calibration. The phosphosites selected for mutation were all selected as having predicted positive effect on the interaction and found near the interface positions of the cognate interaction partner on the AlphaFold predicted structures, as we illustrate for the NKAP-DDX41 interaction (Figure 5A). We confirmed that the mutants do not have reduced expression levels (Figure S2), ruling out changes of interactions by simple loss of protein stability and degradation. The wild-type baits efficiently recovered known interaction partners, with many of them among the top-enriched prey (Figure S3). The 4 replicate experiments run per pull-down correlated strongly across experiments, and correlation coefficients between mutants of the same bait were similarly high (Figure S3), consistent with the expectation that individual phosphosite mutations would cause only small perturbations to the full set of interactions.

**Figure 5.**
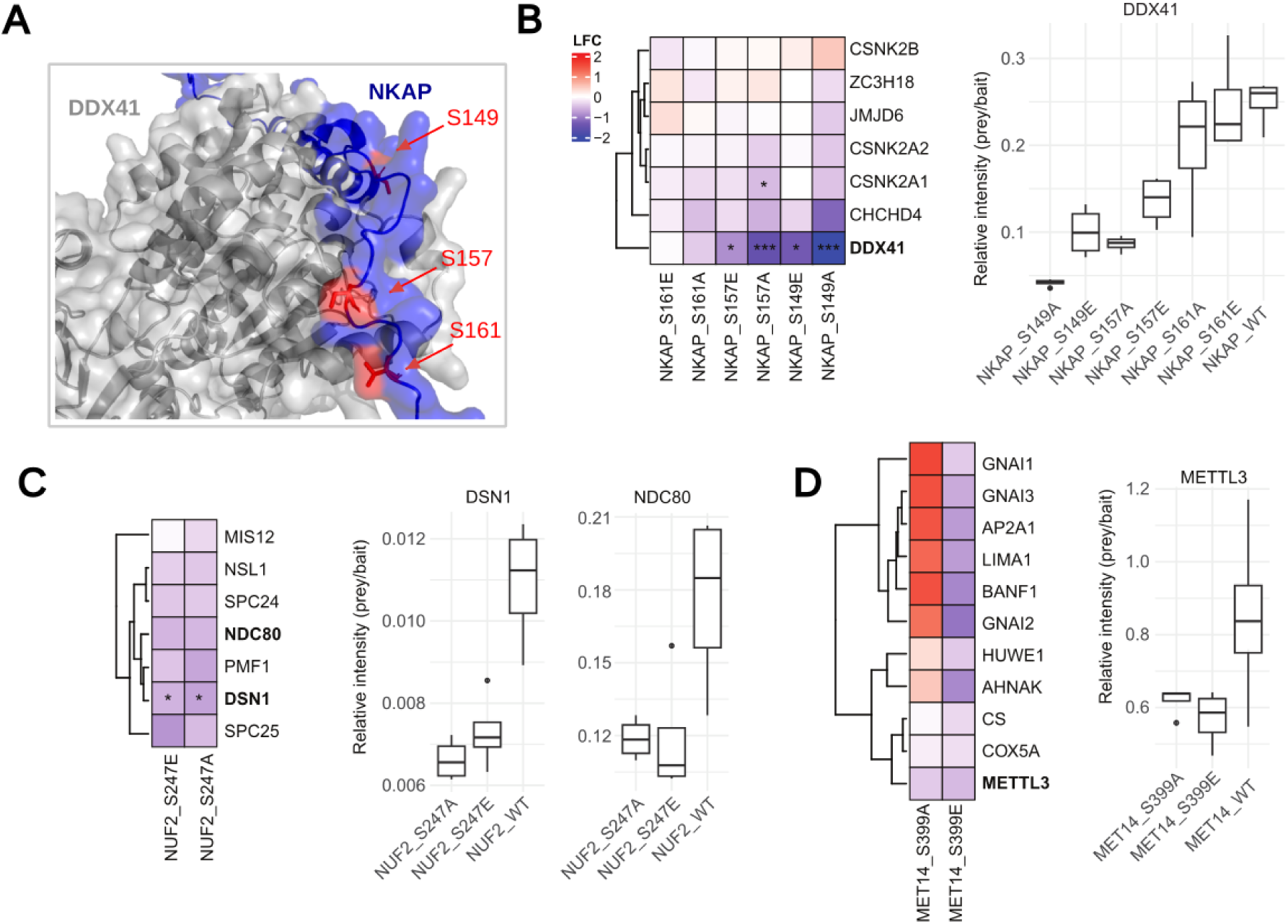
Validating the impact of phosphosite mutations on protein-protein interaction by AP-MS. (A) AlphaFold model for the NKAP-DDX41 interactions, an example of a predicted structure for interactions with putative regulatory phosphosites near the predicted interface that were selected for experimental testing through mutations. (B-D) Quantification in log-fold change (LFC) and relative intensity (prey/bait) of bait-prey interactions from AP-MS for the 3 selected baits NKAP (B), NUF2 (C) and METTL14 (D). Significant changes are denoted in the heatmap (* < 0.05, ** < 0.01, *** < 0.001).

To quantify the effects of the phosphosite mutants, we focused on known interactors curated from BioGRID comparing the abundance of the known interaction partners between the WT and the mutant baits. For NKAP, mutations at S149 and S157 led to significant reductions in DDX41 abundance, while other known interactors remained largely unaffected (Figure 5B). Mutation of S161 did not alter protein interactions of NKAP. Inspecting the predicted structural model, we can note that S149 is buried at the interface while S157 and S161 are near but not buried in the interface. For NUF2, all known interactors showed a slight decrease in intensity, with only a significant decrease of DSN1 (Figure 5C). As NUF2 is part of a large complex together with these proteins (Yatskevich et al. 2024), partial destabilization of the complex could explain this consistent decrease. For METTL14, the positive control phosphosite mutation resulted in a more specific reduction of METTL3 intensities, although this trend was not significant (Figure 5C). Across all examples, we did not observe a difference when comparing the alanine and phosphomimetic mutations. Our initial expectation would have been that the phosphomimetic would promote the interaction, relative to the alanine mutation. However, it is also commonly known that the mimetic mutations don’t often truly mimic the effect of a phosphosite. Overall, 4 out of 5 predicted effects showed consistent decrease in binding relative to the WT but only 2 were statistically significant. Given that the A and E mutants often behaved in a similar manner we also tried combining them to gain statistical power. When doing the analysis in this way the NUF2 S247 mutants show a statistical significant reduction in SPC25 (*p-value* = 0.027) and PMF1 (*p-value* = 0.012). Overall, these experiments show that this framework can discover novel interaction-regulatory sites while also cautioning that there will be a degree of false-positive predictions.

## Discussion

In this study, we present a systematic, multi-omic framework for identifying phosphorylation sites that regulate protein abundance through modulation of protein-protein interactions across the human proteome. By applying co-abundance modeling to pan-cancer CPTAC data encompassing 1,006 tumor samples, we identified 3,038 phosphosites predicted to modulate 6,160 phosphosite-protein associations. While trying to control for transcriptional and clinical confounders, and trying to be stringent by requiring the associations to be present at copy-number and transcriptional level, such associations should be considered as hypothesis generation requiring further testing for validating.

It is worth being explicit about the scope of the causal reasoning that we described in the results section. The requirement for consistent copy-number and transcript effects constrains the direction of the protein-protein relationship, supporting the interpretation that the abundance of protein X shapes that of its partner Y rather than the reverse. This directional argument does not, however, necessarily transfer directly to the phosphosite layer. An association between a phosphosite on X and the abundance of Y remains correlational although the prior protein-protein analysis makes the causal directionality more likely.

In an attempt to give more credibility to the associations we mapped the putative regulatory phosphosites to structural models of protein complexes and to prior proteomics experiments identifying phosphosites with a role in protein localization. We identified 239 phosphosites near interaction interfaces, a number that might be limited still by the incomplete structural coverage of all possible interface regions. Part of these limitations are due to limitations in AlphaFold itself that is, for example, biased towards predicting certain types of interactions better than others (Burke et al. 2023). We note that the cut-off used (<8A based on all-atom distance) is lenient and some of these positions might not contact the interface but be near the interface. Beyond interface regulation, our findings implicate subcellular localization as an equally important dimension of phosphorylation-mediated interaction control. Among our predicted phosphosites, 402 phosphosites have been previously linked with differences in subcellular localization showing how this might affect complex formation and partner stability in a compartment-specific manner.

There are several limitations of our study worth noting. The reliance on predicted rather than experimentally determined complex structures introduces uncertainty, particularly for phosphosites within intrinsically disordered regions where structural confidence scores are inherently lower. Similarly, the pan-cancer context of the CPTAC dataset, while providing statistical power, conflates regulatory phosphorylation events across diverse tumor types and may obscure cancer-type-specific mechanisms. Future studies applying this framework to individual cancer types or non-malignant tissues may reveal additional context-dependent regulatory relationships. The majority of the phosphosites that we predict to regulate interactions were not assigned to interfaces or regulation of localization which we believe is primarily due still to the limitations in these two annotation sources. There are however other mechanisms by which phosphorylation of a protein would alter protein interactions indirectly, including for example by regulating conformational changes (Correa Marrero et al. 2025). Finally, the experimental validation, while done on a relatively small number, already showed a degree of likely false positive predictions. The list of associations can be best seen as a way to enrich for the most plausible candidates to further study experimentally.

Taken together, this work establishes co-abundance modeling integrated with structural predictions as a powerful and scalable approach for functionally prioritizing phosphorylation sites across the proteome. The resulting resource comprising thousands of phosphosite-protein associations annotated with structural, localization, and functional information is provided in supplementary information (Table S1) for targeted experimental follow-up. We think this can contribute towards advancing our understanding of how post-translational modifications regulate protein interaction networks.

## Materials and methods

### Multi-omics Data Collection

Transcriptomics, proteomics, and phosphoproteomics quantifications at the protein and phosphosite levels were obtained from the CPTAC data portal (Li et al. 2023) using the Python API with the Baylor College of Medicine (BCM) pipeline. For the phosphoproteome dataset, we used a modified API to extract the original data without aggregating protein ensembl IDs. Clinical metadata for all samples was also obtained using the Python API. Copy number variation (CNV) GISTIC 2.0 levels were compiled from the LinkedOmics portal (https://www.linkedomics.org/, accessed July 12, 2024), where discrete variables of -2, -1, 0, 1, and 2 are recorded.

### Data Preprocessing and Normalization

Non-tumor samples were first removed from the dataset. For the phosphoproteome data, identical phosphosites were aggregated by taking the median values. Transcript, protein, and phosphosite quantifications were then normalized for each protein or phosphosite by subtracting the median log2 intensity across all samples from individual log2 intensity values. Clinical metadata was linked to each sample, and only samples with complete age and sex information were retained for subsequent analysis. The age with ‘>90’ was converted to integer 90. The final dataset comprised 1,006 samples with CNV, transcriptome, proteome, phosphoproteome, and metadata information. Due to the sparse nature of proteomics and phosphoproteomics data, we applied stringent filtering criteria: proteins were required to be measured in at least 200 of the 1,006 samples, yielding 11,231 proteins for analysis. Similarly, phosphosites measured in at least 200 samples were retained, resulting in 28,318 phosphosites across 6,635 proteins.

### Compendium of Physical Protein Interactions

We constructed a comprehensive catalog of physical protein interactions using two complementary approaches as described previously (Sousa et al. 2019). First, we downloaded protein complex data from the CORUM database (accessed July 16, 2024) (Tsitsiridis et al. 2023; Steinkamp et al. 2025), reasoning that proteins within the same complex interact physically at least once. From 3,694 protein complexes, excluding homodimers, we assembled 91,754 protein interactions (45,877 unique pairs). Second, we obtained protein-protein interaction data from BioGRID version 4.4.235 (Oughtred et al. 2021), selecting only human interactions captured through physical experimental systems. We excluded Affinity Capture-RNA and Protein-RNA interactions to ensure our dataset contained only protein-level interactions. After removing homodimers, this yielded 1,641,206 protein interactions (820,603 unique pairs). Combining CORUM and BioGRID data, we established a comprehensive compendium of 1,686,898 protein interactions (843,449 unique pairs).

### Linear Modeling to Identify Protein Associations

For a given protein physical interaction pair X and Y, it was tested whether protein X can control the protein levels of Y through protein–protein interaction, potentially constraining the degradation rate of Y. For each interacting pair, two nested linear models were fitted the same as described previously (Sousa et al. 2019). The first model (null) was used to predict the protein levels of Y (Py) using its mRNA (Ty) and a set of other covariates, i.e. cancer type, patient age, and gender (Equation (1)). In a second linear model (alternative), the CNV levels of X (Gx) were added as a predictor variable (Equation (2)). A likelihood ratio test (LRT) (Equation (3)) was then applied to test whether the second model increases the goodness of fit of the first model in predicting Py.

For each protein interaction pair (X and Y), we tested whether protein X could regulate the protein levels of Y through protein-protein interactions, potentially by constraining Y’s degradation rate. We employed nested linear regression models to control confounding factors and isolate the specific effects of interest.

We fitted two models for each interacting pair. The null model predicted protein levels of Y (Py) using its corresponding mRNA levels (Ty) and covariates including cancer type, patient age, and gender:

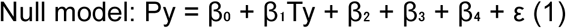

where β_0_ represents the intercept, β_1_ the regression coefficient for mRNA of Y, β_2−4_ the coefficients for cancer type, age, and gender respectively, and ε the error term.

The alternative model incorporated the CNV levels of protein X (Gx) as an additional predictor:

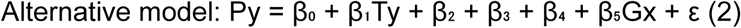

where β_5_ represents the regression coefficient for CNV of protein X.

Model comparison was performed using a likelihood ratio test (LRT):

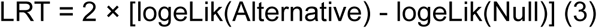

P-values were calculated using the LRT statistic over a chi-squared distribution and adjusted for multiple testing using the Benjamini-Hochberg false discovery rate (FDR) method . This analysis was applied to protein pairs where CNV x, Protein y, and Transcript y data were available, encompassing 1,099,864 protein associations across 1,006 tumor samples. We also performed parallel analyses substituting mRNA levels for CNV of protein X, by substituting the term Gx to the transcript levels of protein X (Tx).

### Phospho–protein Associations

To investigate whether phosphosites on protein X could influence the protein abundance of interacting partner Y, we employed an extended modeling approach as described previously. The null model predicted protein levels of Y (Py) using its mRNA (Ty), the CNV and protein levels of X (Gx and Px), and the same covariates:

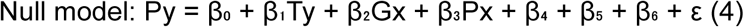

The alternative model incorporated the phosphosite levels of protein X (Px^ph^):

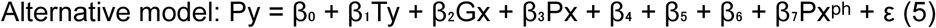

where β₇ represents the regression coefficient for the phosphosite of protein X. Models were compared using the same LRT framework described above.

This analysis was applied to phosphosite-protein pairs where all required variables (Px^ph^, Px, Gx, Ty, and Py) were available, resulting in 4,106,384 tested pairs across 1,006 tumor samples.

### Subcellular localization Analysis

To identify phosphosites that regulate protein subcellular localization among those showing significant associations, we employed a two-pronged approach combining database mining and literature curation.

First, we accessed the PhosphoSitePlus repository and downloaded the Regulatory Sites table (accessed December 18, 2024). Phosphosites annotated with “intracellular localization” function were classified as having regulatory roles in protein localization. Second, we reviewed published literature to identify phosphosites that exhibit localization patterns distinct from their parent proteins. We compiled data from four complementary studies (Masuda et al. 2020; Xu et al. 2023; Krahmer et al. 2018; Martinez-Val et al. 2021) to construct a comprehensive catalog of phosphosites with altered subcellular distribution profiles. We directly adopted their experimentally validated phosphosites from Masuda et al. and Xu et al. Similarly, from Krahmer et al., we adopted documented phosphosites after converting mouse phosphorylation sites to human orthologs using SITE_GRP_ID identifiers from the PhosphoSitePlus database. For Martinez-Val et al., which provided subcellular fractionation profiles without explicit phosphosite annotations, we applied the analytical workflow described by Masuda et al. to identify relevant phosphosites. Specifically, both phosphosite and protein abundance values were normalized to a 0-1 scale across six subcellular fractions. Phosphosites were considered to have distinct localization patterns if they exhibited significantly different enrichment ratios compared to their parent proteins (p < 0.01, with maximum difference in enrichment ratios > 25% in any fraction).

### Structural Analysis

We used previously predicted protein-protein interaction models from (Jänes et al. 2024). Interactions not covered by these models were predicted using ColabFold v1.5.5 (Mirdita et al. 2022) with --num-models 1.

### Plasmid construction and protein expression

The cDNAs encoding METTL14, NKAP, and NUF2 were obtained from pDONR223 vectors selected from the human Gateway ORFeome collection. NKAP contained two mutations, G6V and D414H, which were corrected by site-directed mutagenesis PCR using primer pairs NKAP-V6G-fwd/NKAP-V6G-rev and NKAP-H414D-fwd/NKAP-H414D-rev, respectively. To generate SH-tagged bait proteins, the respective cDNAs were introduced into the destination vector pcDNA5/FRT/TO/SH/GW (pTOSH-N) (Glatter et al. 2009) using the Gateway™ LR Clonase™ II Enzyme Kit (Invitrogen, 11791020). The ligation mixture was transformed into chemically competent E. coli DH5α (NEB, C2987H). Phosphomimetic (Glutamate, E) and alanine mutants were generated using Phusion HF polymerase (NEB, M0530S) and the Q5® Site-Directed Mutagenesis Kit (NEB, E0552), with two pairs of primers used for each site (see list of primers). All constructs were confirmed by full plasmid sequencing (Microsynth AG, Switzerland). HEK293T cells were used for protein expression, following an adapted protocol from (Liu et al. 2020; Kaushal et al. 2024). Briefly, cells were cultured in 10 mL DMEM high glucose pyruvate (Thermofisher, 41966029), supplemented with 10% fetal bovine serum FBS (Thermofisher, A5256801) and 1% Penicillin–Streptomycin (Thermofisher, 15140122), in 10-cm dishes. The next day, when cells reached 80% confluency, each plate was transfected with 15 µg of plasmid DNA following the JetPEI manufacturer’s protocol (Sartorius, 101000053), with four replicates per bait. After two days of culture, cells were washed with 5 mL PBS at room temperature and immediately detached with 5 mL PBS pH 7.4 (1X) [Thermofisher, 10010023] and collected in 15-mL Falcon tubes on ice and centrifuged for 5 minutes at 1200 rpm, 4 °C. The supernatant was discarded, and the cell pellets were quickly shock-frozen in liquid nitrogen and stored at -80 °C until further use.

### Affinity purification of the interacting proteins

Cells were resuspended in 1 mL of ice-cold HENN lysis buffer (50 mM HEPES, 150 mM NaCl, 1 mM EDTA, 50 mM NaF, 0.4 mM NaVO_3_; pH 7.5) supplemented with 0.5% IPEGAL CA-630, 1 mM DTT, 0.5 mM PMSF, protease inhibitor cocktails (Roche, 05056489001), and phosphatase inhibitors (Roche, 04906837001). Cells were lysed by gentle vortexing for 5 sec, incubating on ice for 5 min, and pipetting up and down, followed by centrifugation at 13,000 × g for 20 min at 4°C. The cleared lysate was transferred into 2 mL Eppendorf Protein LoBind tubes containing 30 μL of pre-equilibrated MagStrep Strep-Tactin™ XT beads (IBA Lifesciences, 2-5090-010). Tubes were rotated for 2 h at 4 °C. Beads were collected on a magnetic rack (Thermofisher, 12321D) and washed twice with 1 ml HENN wash buffer (50 mM HEPES, 150 mM NaCl, 1 mM EDTA, 25 mM NaF, 0.5% IPEGAL CA-630, 1 mM DTT), followed by three washes with detergent-free HEN buffer (50 mM HEPES, 150 mM NaCl, 1 mM EDTA,1 mM DTT). All the above steps were performed in the cold. Beads were resuspended in 100 μl of denaturation-reduction buffer (2 M urea, 50 mM Tris-HCl, pH 8.0, 5 mM DTT) and incubated at 37 °C, 700 rpm for 30 min. Proteins were then alkylated with 15 mM iodoacetamide (IAA) at room temperature in the dark, followed by quenching with 10 mM DTT for 15 min at room temperature. This was immediately followed by on-bead tryptic digestion.

### Tryptic digestion and peptide C18-cleanup

On-bead digestion was performed by adding in 1 µg trypsin (Promega, V5113) into the tubes that contain the beads, and the reaction was carried at 37 °C overnight at 700 rpm on a thermomixer. After digestion, beads were collected on a magnetic rack, and the supernatant was transferred to a 1.5 mL Protein LoBind tube. Beads were resuspended in 50 µL of 50 mM Tris-HCl (pH 8.0), and the eluate was pooled with the first supernatant. The resulting tryptic peptides were acidified with TFA to a final concentration of 1.5% before peptide clean up. Peptides were purified using a µHLB Oasis 96-well plate (Waters, SKU: 186001828BA) connected to a vacuum pump, following the manufacturer’s recommendations. Briefly, the plate was conditioned sequentially with 200 µL methanol, 200 µL of 40% acetonitrile (ACN) containing 0.5% acetic acid (AA), and three washes with 200 µL of 0.1% trifluoroacetic acid (TFA) in water. Acidified tryptic digest samples were loaded onto the plate and washed three times with 200 µL of 0.1% TFA in water, under gentle vacuum pump pressure. Peptides were eluted in two steps: first with 80 µL of 40% ACN containing 0.5% AA, then with 100 µL of 70% ACN containing 0.5% AA. Eluted peptides were dried in a SpeedVac at 46 °C and stored at -80 °C until further use.

### LC-MS analysis

DIA-MS measurements were performed on an Orbitrap Astral mass spectrometer coupled to a Vanquish Neo UHPLC system (Thermo Fisher Scientific). Dried peptides obtained as described above were resuspended in 20 µL of 5% acetonitrile (ACN) with 0.5% formic acid (FA). A 0.5 µL aliquot of the peptide solution was loaded by direct injection and separated onto a 75µm diameter nano emitter (CoAnn) self-packed with 25cm of 3µm ReproSil-Pur C18-AQ particles (Dr Maisch), maintained at 50 °C. Peptides were separated using a binary solvent system consisting of solvent A (water with 0.1% FA, v/v) and solvent B (80% ACN with 0.1% FA, v/v). The peptides were loaded using the fast loading combined-control mode limited to 1µLl/min or 800nl/min. The gradient started with a linear increase from 3% to 32% B over 15 min while the flow rate decreased from 800nl/min to 400nl/min. The gradient then increased to 50% B for 5 min at 4 00nL/min, followed by a ramp to 95% B within 1 min and column flushing at 95% B for 5 min. Finally, the column was re-equilibrated to 5% B over 2 min at 800nl/min, giving a total run time of 28 min. For mass spectrometry analysis, data-independent acquisition (DIA) was performed in positive ion mode with an LC peak width set to 10 and a default charge state of 2. MS1 and global settings were as follows: ion transfer tube temperature, 275 °C; spray voltage, 2.5 kV (controlled by the Tune settings); RF lens, 40%; and full MS1 resolution, 240,000. The AGC target was custom (normalized AGC target = 500%), with a maximum injection time of 3 ms, one microscan, and a scan range of 350–1400 m/z. DIA-MS2 spectra were acquired with a normalized AGC target = 500%, with one microscan and isolation windows of 2 m/z (524 scan events) with the window placement optimization option activated. The precursor mass range was set to 350–1400 m/z, and fragment ions were scanned over a range of 150–2000 m/z. Higher-energy collisional dissociation (HCD) was applied with a normalized collision energy (NCE) of 25%.

### Affinity Purification Mass Spectrometry Data Analysis

To identify high-confidence interactors for wild-type and phosphosite mutant NKAP, METTL14, and NUF2, we searched the samples separately per condition using directDIA™ in Spectronaut™ (v. 20), following the standard pipeline. Searching each condition independently prevented peptides from being identified in one condition based on detections in another. We filtered for proteins with a Q value < 0.05 and with at least two unique peptides. We then log2 transformed the intensity values and normalized the data with median-balanced quantile normalization. Then, we input all samples into SAINTexpress v3.6.3 (Teo et al. 2014) and calculated the confidence scores of the protein-protein interactions. For visualization, we filtered the preys with Bayesian false discovery rate (BFDR) ≤ 0.001, Saint Score of 1, fold change from control > 2, and average intensity across replicates > 2. We also downloaded all interactors per bait from BioGRID (https://thebiogrid.org/) and downloaded known protein complexes from CORUM (https://mips.helmholtz-muenchen.de/corum/). For further analysis, we only kept preys with BFDR ≤ 0.001 and Saint Score of 1.

For quantification, we re-searched the data using directDIA™ in Spectronaut™ per bait instead of per condition. The same filtering and normalization were applied as above. We calculated Pearson correlation coefficients between replicates and conditions for every bait, using only proteins that passed the quality control from the initial search. We further filtered the proteins to include only known BioGRID interactors that were identified in the wild-type bait samples with an average intensity across replicates above 1. We calculated log2 fold changes between phosphosite mutants and wild-type baits and tested significance using pairwise two-sided t-tests. We applied FDR correction per bait.

## Acknowledgements

We thank Dr. Peter Francis Doubleday for providing HEK293T cells and helping in the initial protein expression tests. We are grateful to Dr. Matthias Gstaiger for providing the pDONR223 and pTOSH-N plasmids and for sharing his expertise on AP–MS. We also thank the laboratory of Prof. Paola Picotti, in particular Dr. Ludovic Gillet, Dr. Anna Pagotto, and Dr. Alexander Leitner, for their valuable input, MS method development, and for maintaining the Orbitrap Astral instrument. K.O. was supported by JST ACT-X (JPMJAX2324), JSPS KAKENHI (25K18607), and the FY 2024 Researcher Exchange Program between JSPS and ETH. P.B. is supported by the Helmut Horten Stiftung and the ETH Zurich Foundation.

## Supplementary Figures and Tables

**Fig S1.**
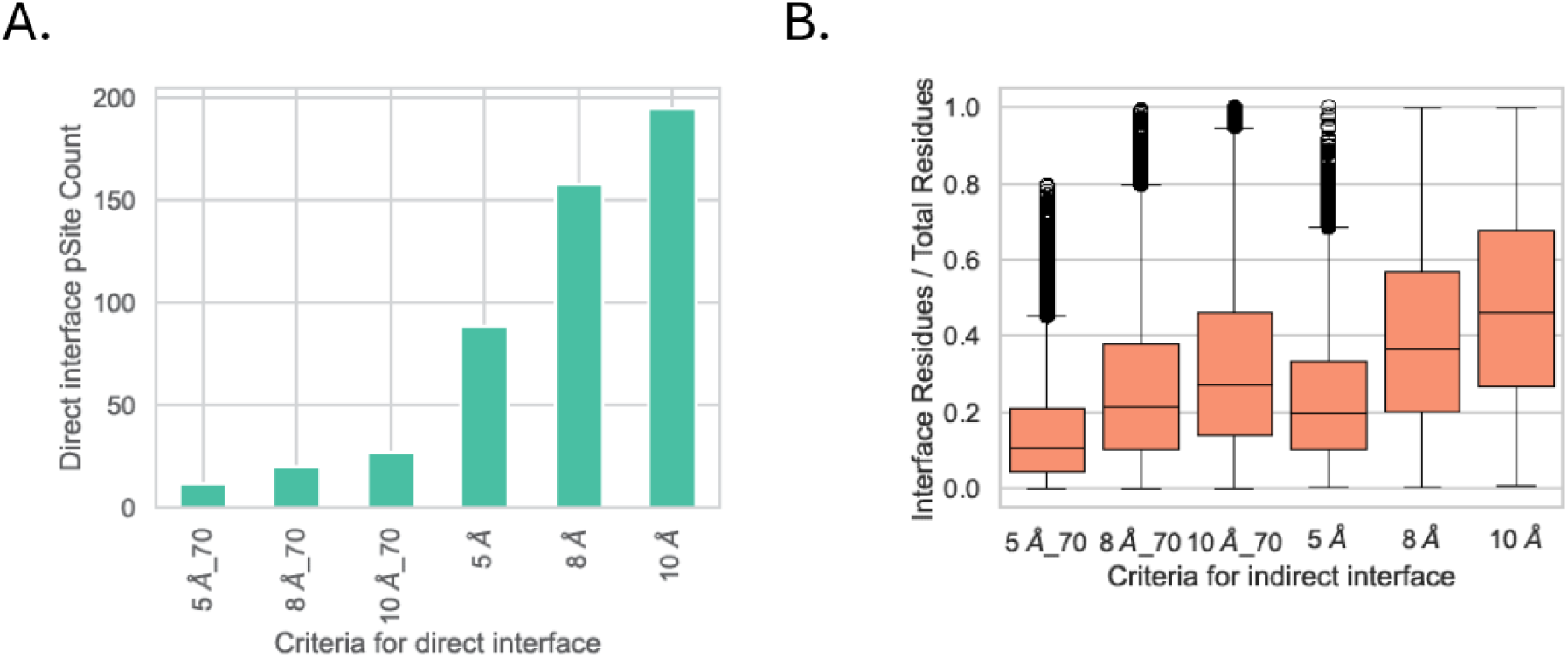
Classification of interface residues and phosphosites. Predicted heterodimer structures were used to identify amino acid residues at the interface. Interface residues are defined by their proximity to the nearest residues on interacting proteins. (A) Interface phosphosites were categorized using predicted structure of specific pairs derived from regression analysis with various distance cutoff values. “_70” indicates that only residues with pLDDT > 70 are considered. (B) Interface residues, irrespective of the interacting proteins on a specific protein, are identified using all available heterodimer structures with pDockQ > 0.5. Relaxed criteria sometimes categorize all residues in the protein as interface residues, indicating over-annotation.

**Fig S2.**
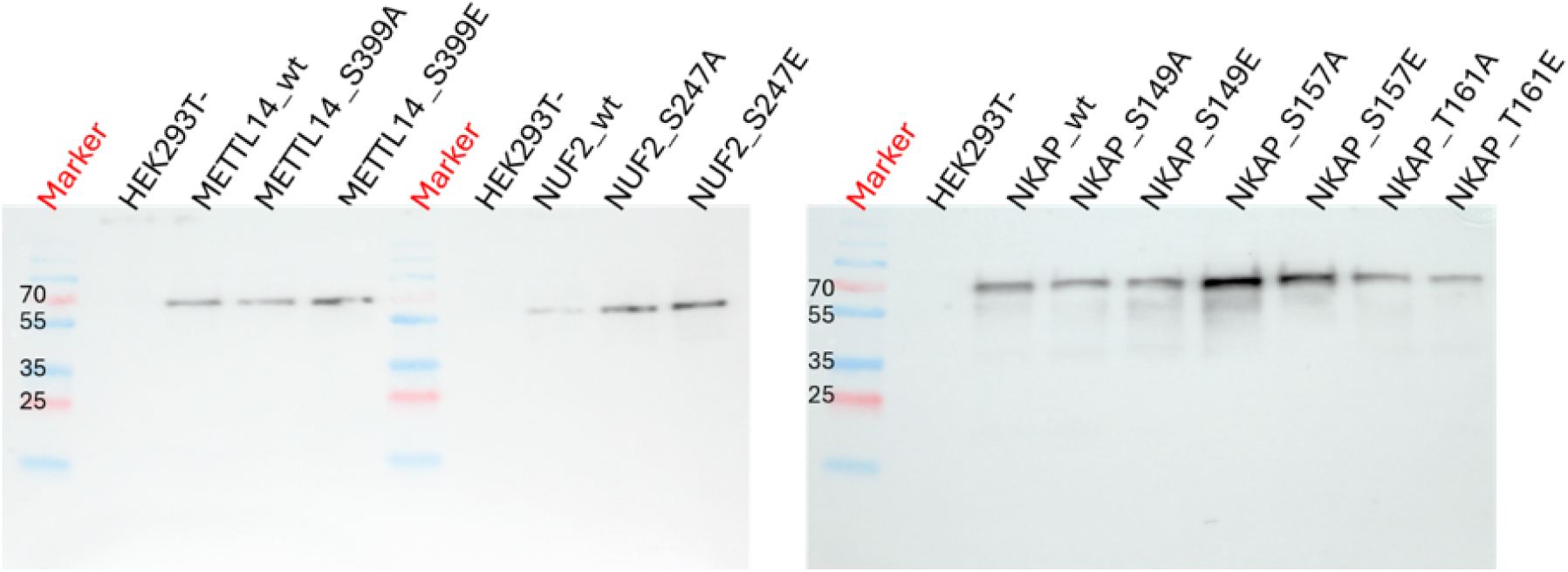
Western Blot analysis of expressed tagged proteins and mutants. Protein expression in HEK293T and cell lysis were performed as described in the Materials and Methods section. 50 µL of cell lysate was diluted with 4x Laemmli sample buffer (Bio-Rad, 1610747) and boiled at 95 °C for 5 min. 15µL of the solution was loaded on a 10-well 10% gel (Bio-Rad, 4569033EDU). Strep II tagged proteins were detected using mouse Anti-Strep antibody (Qiagen, 34850), and Goat anti-Mouse IgG, HRP-linked Antibody (Invitrogen, A16078) as the primary antibody and secondary antibody, respectively. Proteins were visualized using Fusion FX (Vilber). The marker is PageRuler™ Prestained Protein Ladder (Thermo Scientific, 26616).

**Fig S3.**
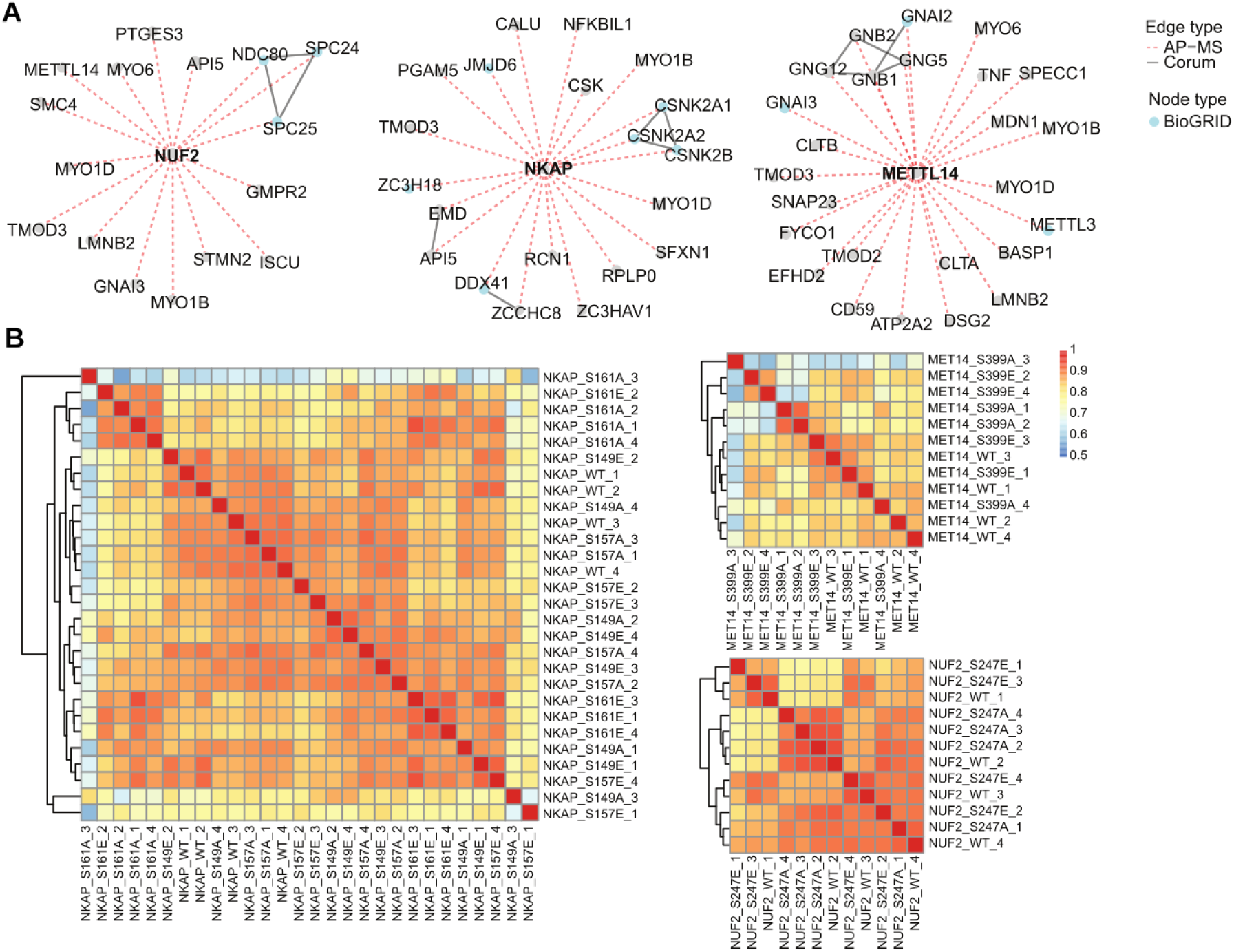
AP-MS of selected mutant bait proteins. We selected bait proteins with predicted regulatory phosphosites (NUF2, NKAP and METTL14), generating mutants for the phosphosite positions predicted to be regulatory. (A) Top identified proteins from the pull-down experiments of the WT proteins have many known interactors found in the BioGrid database (blue dots). (B) The heatmaps show the correlation of intensities of the prey proteins identified for replicates of the different mutants showing an overall very high correlation value.

## List of supplementary files

Table S1 - List of significant phospho - protein association pairs. List of significant associations between phosphorylation sites and putative regulated target protein, annotated with information on protein structural interfaces and localization regulation information.

## Notes

### Competing Interest Statement

The authors have declared no competing interest.

